# Metagenomic insight into taxonomic composition, environmental filtering, and functional redundancy for shaping worldwide modern microbial mats

**DOI:** 10.1101/2022.08.09.503407

**Authors:** M. Viladomat, M. García-Ulloa, I. Zapata-Peñasco, L. E Eguiarte, V. Souza

## Abstract

Although microbial mats are considered relictual communities that are nowadays mostly constrained in their distribution by predation and phosphorus availability, they are still found in a wide range of environmental conditions. Their ancestral history, geographical isolation, stratified community composition and interspecies dynamics make them an interesting model to study community ecological processes and concepts. In this study, we analyzed different metagenomic datasets from worldwide modern microbial mats to compare community structure and functions. We found significant differentiation in both alpha and beta diversity of taxonomic and functional categories without significant correlation with temperature and pH. Differences depended more on the presence of very highly abundant cyanobacteria and some generalist microorganisms. Our results suggest that there is more than just Grinnellian niche dynamics in the determination of microbial mat community assembly, opening the hypothesis of interactions as the driver behind these ancient communities. We also discuss the influence of niche dynamics and environmental filtering in the community assembly of microbial mats.

## Introduction

Microbial mats are multilayered communities considered the most primitive communities on Earth, with fossilized material as old as 3.7 billion years [1]. The typical multilayered structure of microbial mats (‘vertical stratification’) originates from physicochemical gradients such as light, oxygen and sulfur that are generated and maintained by the metabolic activity of their members [2]. Physicochemical gradients provide microenvironments for different functional guilds [3] with particular metabolic needs and tolerances [4]. From bottom to uppermost layer, the formation often starts with the strictly anaerobic methanogens, followed by the diverse sulfur bacteria and archaea, then the early phototrophic guilds of purple and green sulfur and non-sulfur bacteria, and finally, cyanobacteria and eukaryotes [2, 5], containing all the known biogeochemical cycles within only a few millimeters [6, 7].

Although microbial mats are thought to have been very abundant in the distant past [8, 9, 10], modern microbial mats are only found isolated from each other, in locations where environmental conditions are often extreme and most grazing eukaryotic growth is limited [11, 12]. Examples are the extremely oligotrophic and hypersaline ponds from the Cuatro Cienegas Basin [13, 14], the high temperature sites from Little Hot Creek and Yellowstone [15, 16, 17] and even on uranium and gold mines [18]. Furthermore, microbial mats have been found to be highly metabolically interdependent and resilient communities [19, 20], where codependence between microbial groups is favored to sustain the entire community [21, 22, 23].

Aside from the Bolhuis et al (2014)[2] mini-review and Prieto-Barajas et al (2018)[24] review on microbial mats at a global scale, the vast majority of studies to date focus on particular sites (site characterization, time series and environmental change experiments) [25, 26, 27, 28, 29, 30] or comparison between similar environmental conditions [31, 32]. These local studies have shown relevant information about the ecology of microbial mats such as their remarkable nutrient scavenging and recycling abilities [32] and their robustness and capacity to endure perturbations due to their functional redundancy [25, 27, 28]. Cardoso et al (2017) [26] found that Cyanobacteria dominated for almost 95% of active microbial mat communities and represented 60% of the resident fraction. Prieto-Barajas et al (2018) [24] described, from bibliographic sources, dominant genera found in different environmental categories, where Cyanobacteria was also found to be the most shared dominant phyla throughout all environmental categories. Nonetheless, Ley et al (2003)[33] found that Cyanobacteria only constituted a large fraction of the biomass in the upper few millimeters (>80% of the total rRNA and photosynthetic pigments), but Chloroflexota sequences were conspicuous throughout the mat. Studies from Arctic and Antarctic protein-coding genes from microbial mats showed similarities between the mats from the two poles, with most genes derived from Cyanobacteria and Proteobacteria [32, 34, 35]. Lastly, Khodadad and Foster (2012)[36], through comparative genomic analyses of the functional genes of non-lithifying and lithifying mats, found extensive similarities in most of the subsystems between the mat types, suggesting a high degree of taxonomic and functional stasis on mats in their functional and cyanobacterial (taxonomic) composition.

Nevertheless, although these particular-site-variation focused studies have given us insight into local and temporal community influences and dynamics, the global comparison of worldwide microbial mats has never been done. We believe that as microbial mats adaptability, robustness and capacity to endure perturbations is explained to come from their functional redundancy, their remarkable nutrient scavenging and recycling abilities, with all these functional cohesion and redundancy we would expect to see a similar functional and very dissimilar taxonomic compositions throughout worldwide microbial mat samples.

To test such hypotheses, in this study, we thoroughly compare the taxonomic and functional composition of modern microbial mats from 8 sites across the globe (63 samples total) using publicly available shotgun metagenomes sequenced through the Illumina platform and submitted as raw reads. Also, we assess the influence of geographic location, temperature and pH on them. Moreover, we describe the sampling, sequencing and assembly of two metagenomes from a newly found hypersaline microbial mat from Cuatro Cienegas Basin that we called Archaean Domes.

## Materials and methods

### Sampling, sequencing and assembly of Archaean Domes metagenomes

Samples were taken under SEMARNAT scientific permit SGPA/DGVS/03121/15 from a recently discovered pond named “Archaean Domes” in Rancho Pozas Azules, located at the southeast of the Sierra de San Marcos within Cuatro Ciénegas Basin (CCB, 26º 49’ 41.7’’ N 102º 01’ 28.7’’ W).

Sampling took place during 2016-2017 at the following times: April 10^th^ 2016 (dry season) and September 3^rd^ 2016 (wet season). Water physicochemical conditions were measured 5 cm deep during the second sampling period, using a Hydrolab MS5 Water multiparameter probe (OTT Hydromet GmbH, Germany) (S2 Table). During the first sampling season the ponds were dry, so no measurements could be taken. From each sampling time, 5 cm^3^ of microbial mat samples were obtained using a sterilized scalpel and transferred to sterile Falcon tubes (50 mL) and immediately placed in liquid nitrogen before being stored at -80 °C; this preserved the mat structure until DNA preparation processing. The DNA extraction from the Archaean Domes samples was performed using a column-based modification of the protocol described by Purdy et al. (1996) [37] for shotgun whole-genome sequencing paired-end 2×300 using Illumina Mi-Seq at CINVESTAV-LANGEBIO, Irapuato, Mexico. Raw metagenomic reads are available at NCBI through the SRA database accession number PRJNA612690.

Raw data from Archaean Domes was quality-checked using FastQC [38]. Indexed adapters and barcodes were removed, and low quality sequences were discarded with Trimmomatic v0.36 using a sliding window of 4 pb and an average quality-per-base of 25 [39]. The assembly of the trimmed reads was conducted with Megahit v1.1.1 using the option *– presets meta-large* [40].

### Metagenome selection

A total of 63 microbial mat shotgun metagenomes from 8 sites across the globe were selected from the MG-RAST repository (Keegan et al., 2016): Archaean Domes CCB (n of metagenomes = 2); Shark Bay (n=15); Little Hot Creek (n=3); Schiermonnikoog (n=17); Death Valley (n =6); Mono Lake (n = 3); Rottnest Island (n = 10); Kowary mine (n=3); and Zloty Stok mine (n=2) (S1 Table). To achieve a robust comparison, we verified all analyzed metagenomes were sequenced with the Illumina platform and submitted as raw reads. Moreover, sufficiency of sampling effort was assessed with rarefaction curves of species richness against number of reads (S1 Fig). MG-RAST accession numbers and coordinates of all metagenomes used in this study (S1 Table) and the temperatures and pH measurements for each site (S2 Table) were found in corresponding publications. For metagenomes without scientific publication, MG-RAST metadata was used.

### Metagenomic annotation

Metagenomic sequences were preprocessed by using SolexaQA [41] to trim low-quality regions from FASTQ data, dereplicated with a k-mer approach [42] and screened for contaminants with Bowtie [43].

Taxonomical and functional annotations were performed using the Last Common Ancestor algorithm [44] and the RefSeq and KO (Kyoto Encyclopedia of Genes and Genomes Orthology) as reference databases. Matches of >25 nucleotides and >65% similarity to a taxonomic group or a function with an E-value of ≤10^−5^ were considered significant and therefore included in the analysis.

### Data processing and analysis

Abundance tables were filtered for categories with <4 hits and <20% prevalence across samples. In order to make highly variable data comparable, resulting tables were rarefied to minimum library size and further normalized by the Trimmed means of M-values (TMM) method, which has proven highly accurate for shotgun data (Pereira et al., 2018) [45].

Statistical analyses were performed on the Microbiome Analyst platform [46]. Alpha diversity was calculated by both the Shannon and Simpson diversity indexes and significance was calculated through the Kruskall-Wallis test. Beta diversity was calculated from Bray-Curtis dissimilarity matrices and visualized through Non-Metric Multidimensional Scaling (NMDS) plots and Ward’s clustering algorithm. Site differences were also assessed quantitatively by analyses of similarities (ANOSIM). Core analysis was performed with a minimum sample prevalence of 25% and minimum relative abundance of 0.01%. The Linear Discriminant Analysis Effect Size (LEfSe) algorithm was used to detect features with significant differential relative abundance across sites with a P-value cutoff of 0.01 after False Discovery Rate adjustment.

Correlations between beta diversity and 1) geographical distance and 2) environmental measurements of temperature and pH were calculated with the current implementation of the Mantel test from Legendre and Legendre (2012)[47] (9999 permutations, Spearman correlation method). In addition to the previously calculated Bray-Curtis dissimilarity matrices, an Euclidean distance matrix of the environmental measurements and a Harvestine distance matrix of the geographical coordinates of the sites were also calculated with *vegan* and the R package *geosphere* v1.5-10 [48], respectively.

Correlation plots of the Spearman’s rho from the top 100 most abundant taxa and functions were drawn with the R package *corrplot v0*.*84* [49].

## Results

### Description of the sites

We analyzed 63 metagenomes from 8 sites: Archaean Domes (n of metagenomes = 2); Shark Bay (n=15); Little Hot Creek (n=3); Schiermonnikoog (n=17); Death Valley (n =6); Mono Lake (n = 3); Rottnest Island (n = 10); Kowary mine (n=3); and Zloty Stok mine (n=2).

Temperature and pH were independent from each other (Spearman’s rho = -0181, p-value = 0.458). Little Hit Creek was the warmest site (average 77°C ± 4.6 s.d.). Measurements of pH for all sites were close to neutrality or slightly alkaline (6.5 to around 8), except for Archaean Domes which was more alkaline (9.94) and Kowary which was more acidic (average 5.9 ± 0.12 s.d.).

An Euclidean distance matrix of the environmental variables was calculated, and depicted in an NMDS plot (Fig 1). Results showed consistent sample clustering by geographical site, except for Schiermonnikoog (green) and Mono Lake (orange). Furthermore, measurements from Little Hot Creek group separately from the rest as a result of both their high temperature and lower than average pH. Contrastingly, Zloty Stok and Kowary group together, leading us to consider all of their samples under the same category of mines.

**Fig 1.**
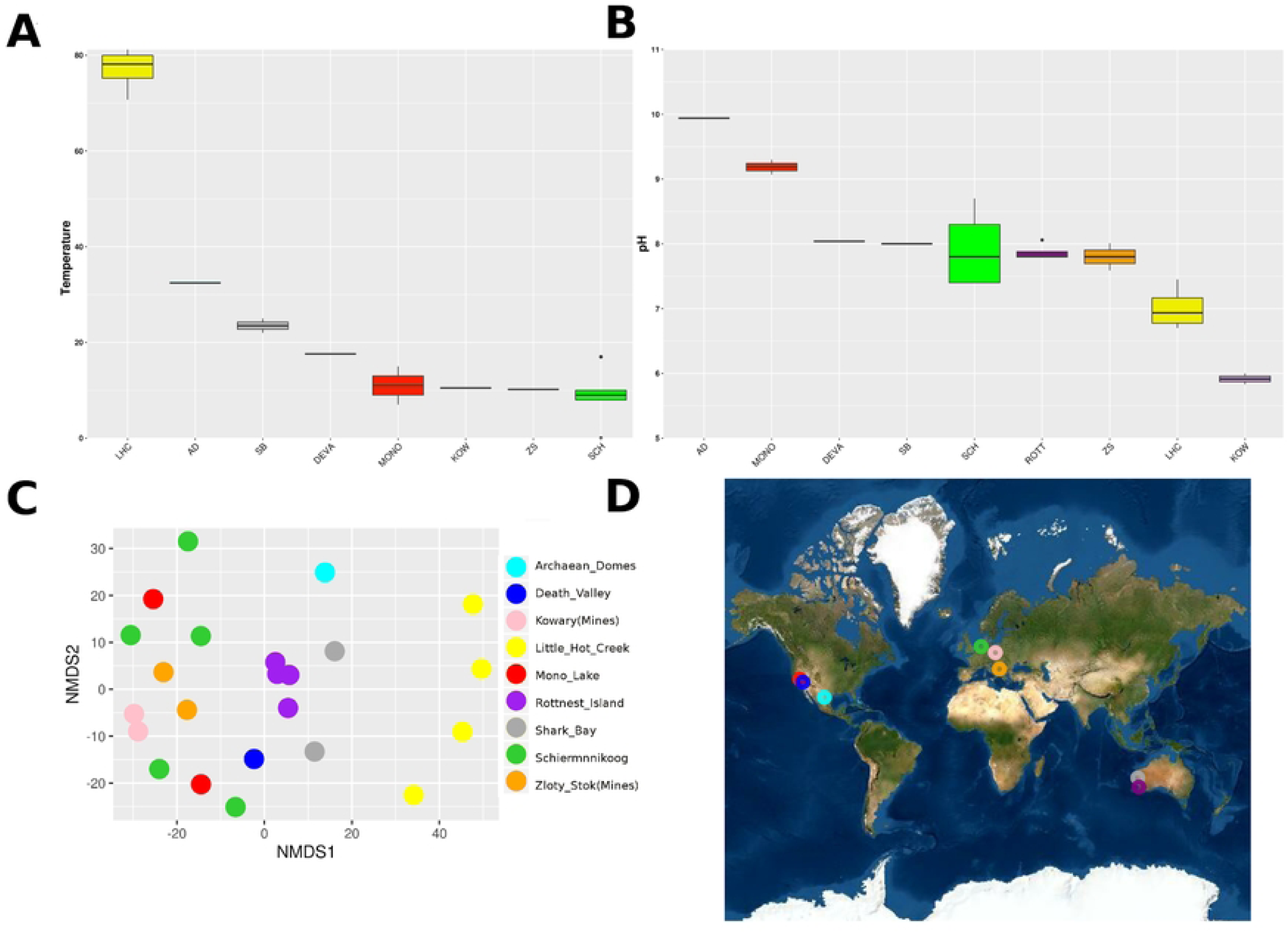
Environmental conditions and geographical location of microbial mats. A) average temperature; B) average pH. Sites are sorted descending according to their mean value. Individual values of environmental measurements can be found in S2 Table. C) NMDS of environmental variables by the Euclidean distance of temperature and pH as a compound variable (ANOSIM’S R = 0.78, p < 0.01; stress score = 0.12). D) Geographic location of each microbial mat. Sites are colored as follows: Archaean Domes (AD, cyan); Death Valley (DEVA, blue); Kowary Mines (KOW, pink); Little Hot Creek (LHC, yellow); Mono Lake (MONO, red); Rottnest Island (ROTT, purple); Shark Bay (SB, gray); Schiermoonnikog (SCH, green); and Zloty Stok mine (ZS, orange). Rottnest is missing from the temperature plot as there were no measurements available.

Shark Bay and Schiermonnikoog as well as Archaean Domes, Death Valley and Mono Lake metagenomes showed a highly similar pattern of variation at the genus level within each site cluster. However, notable differences occurred in the mines cluster, Little Hot Creek and Rottnest: the mines show a general lower diversity and higher abundance of Candidate *Solibacter*; Little Hot Creek also has lower diversity alongside a higher abundance of *Salinibacter, Roseiflexus, Chlorobium, Chloroherpeton* and *Anabaena*; Rottnest had a high abundance of *Salinibacter* and *Rhodothermus*. On the other hand, KEGG metabolism categories appeared very similar across sites and samples, with the slight exception of the KOW_3 sample from the mines group, which has a unique spike in the metabolism of terpenoids and polyketides (Fig 2).

**Fig 2.**
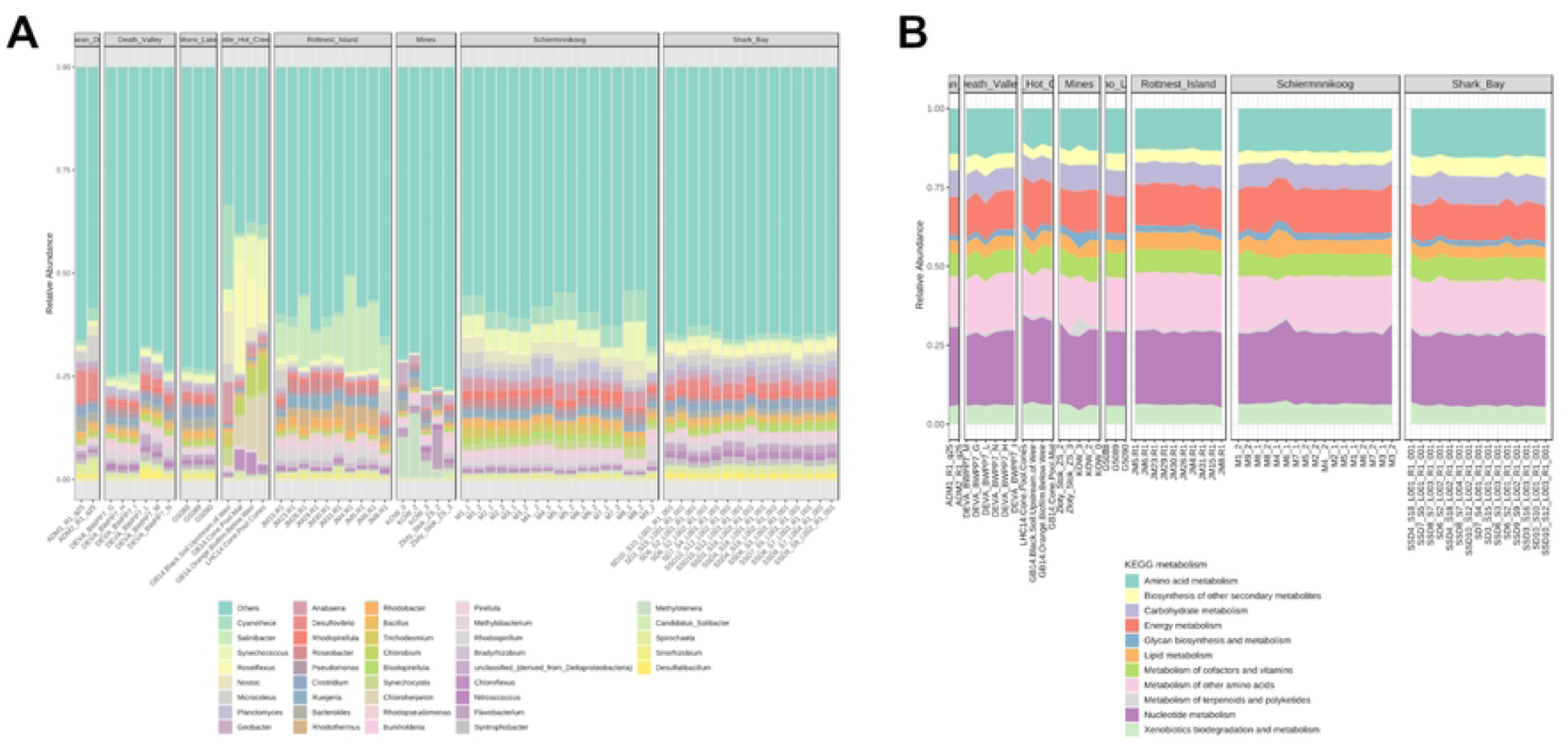
Relative abundance of genus level and function categories of microbial mats. A) The genus stacked bar plot shows the 40 most abundant genera B) while the function shows all available KEGG metabolism categories.

### Taxonomic and functional diversity of microbial mats

Shannon’s diversity of genera is significantly different across sites, ranging between 3.75 and 5.75 (Fig 3. Kruskall-Wallis statistic = 52.0, p = 5.8E-9). Most diverse microbial mats are those from hypersaline Death Valley and Mono Lake sites, closely followed by Shark Bay. Little Hot Creek has the lowest alpha diversity by a significant margin. Contrastingly, functional alpha diversity shows a much narrower range, albeit significantly different across sites, between 4 and 4.18 (Krustall-Wallis statistic = 26.9, p = 3.3E-4), with the exception of KOW_3 with the lowest Shannon value of 3.88. Noteworthy, samples from the mines cluster have both the highest and lowest functional diversity.

**Fig 3.**
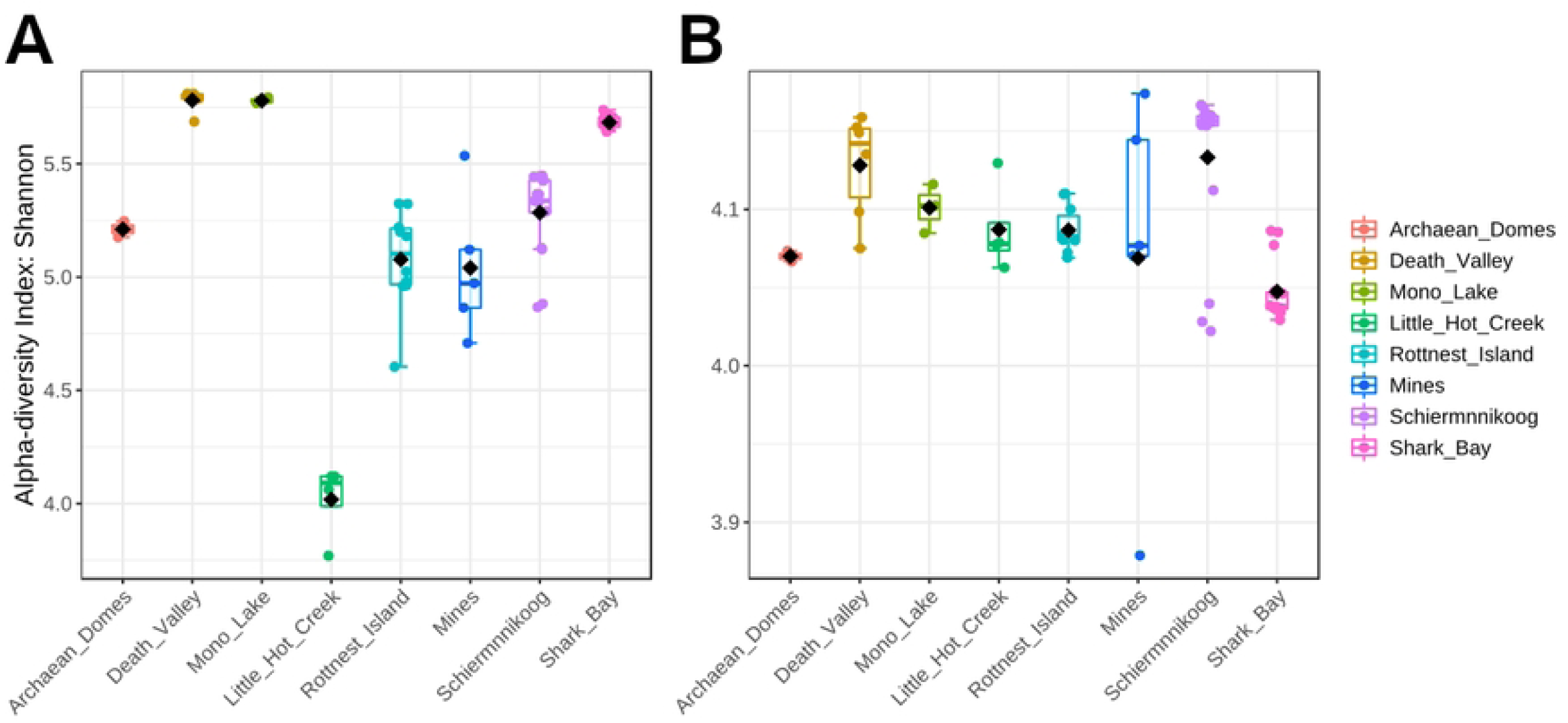
Alpha diversity from microbial mat communities. Alpha diversity at A) genus level and B) KEGG pathways were calculated with Shannon’s Diversity Index. Kruskall-Wallis test gave significant results for both comparisons (statistic = 52.0, p = 5.8E-9 for genus; statistic = 26.9, p = 3.3E-4 for functions).

NMDS plots of beta diversity for genera (Fig 4) show a mostly consistent grouping of samples into their corresponding sites, except for Death Valley which forms two separate clusters. Also, mines and Little Hot Creek samples cluster separately and further from the rest of the sites, displaying low alpha diversity. Contrastingly, clustering of functions is less clear. Most samples from mines, Little Hot Creek, Archean Domes and Schiermonnikoog, as well as some from Rottnest and Shark Bay appear all scattered across. Interestingly, there is a cluster of highly diverse samples and diversity seems to somewhat decrease the further apart the samples are from it, particularly up and to the left as seen by the KOW_3 sample (fully green), some Schiermonnikoog and many Shark Bay samples, leading to a potentially important role of alpha diversity in this ordination analysis.

**Fig 4.**
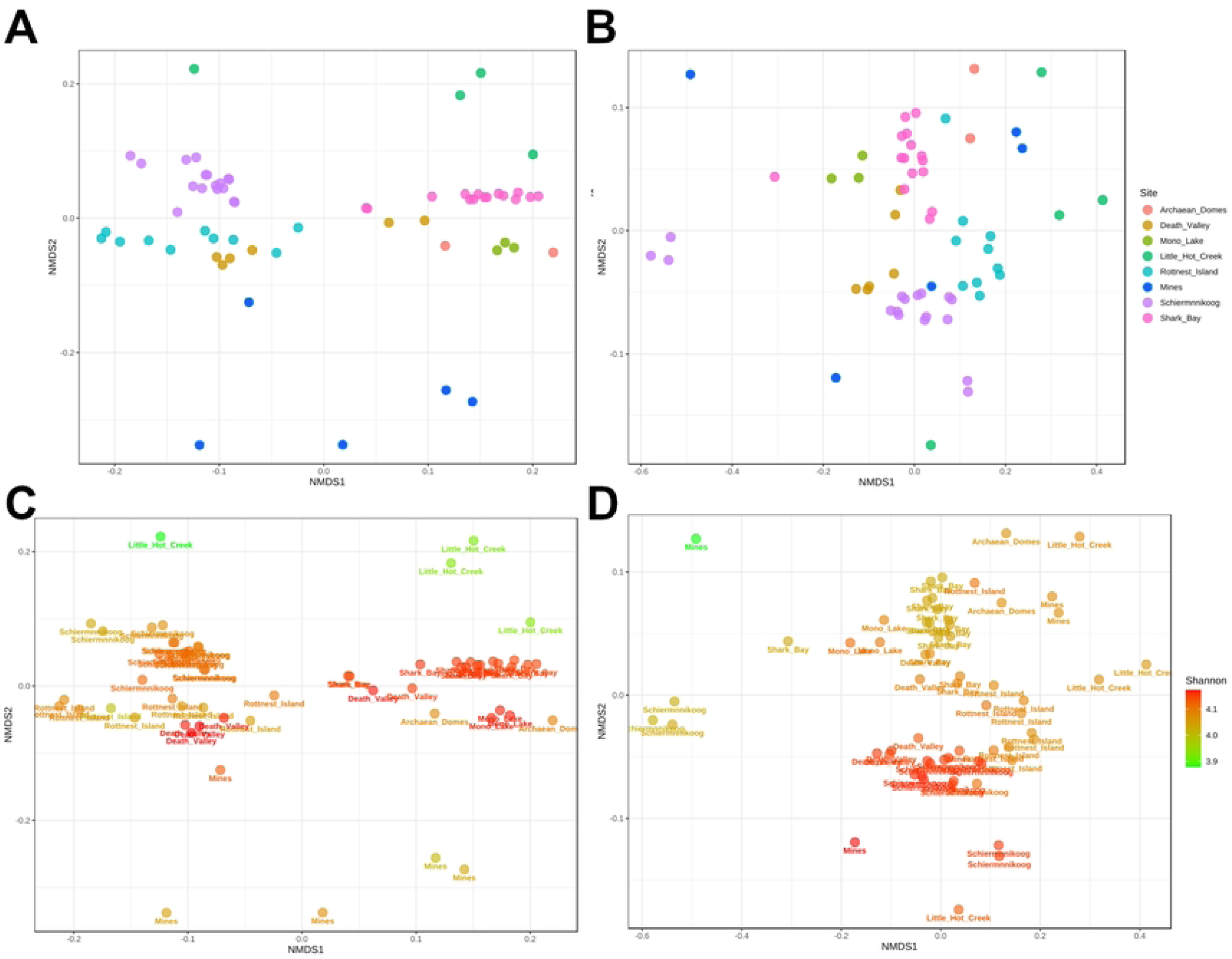
Taxonomic and functional beta diversity of microbial mats. Bray-Curtis dissimilarity was used as a beta diversity metric. A) and C) correspond to genera (ANOSIM’s R = 0.91; p <0.001; stress score = 0.1) and B) and D) to functions (ANOSIM’s R = 0.67; p < 0.001; stress score = 0.08). Bottom panels include Shannon values as a color gradient.

### Similarities and differences across microbial mats

*Cyanothece* is the most abundant and prevalent member of the taxonomic core, with a relative abundance of at least 1% in 71% of the samples, followed by *Microcoleus, Planctomyces* (both 1% relative abundance in around 60% of the samples) and *Rhodopirellula* (1% relative abundance in 50% of the samples) (Fig S2).

Top 40 most significantly abundant genera found with the LEfSe algorithm (Fig 5) include *Clostridium* in Mono Lake; *Planctomyces, Trichodesmium* and *Rhodopirellula* in Schiermonnikoog; multiple sulfur-related *Desulfovibrio, Desulfonatronospira, Desulfohalobium* and *Desulfomicrobium* alongside *Halanaerobium, Spirochaeta* and *Bacteroides* in Archaean Domes; red-pigmented and halophilic bacteria *Rhodothermus, Rhodobacter, Rhodomicrobium* and *Roseobacter, Salinibacter* and *Ruegeria* alongside *Methylobacterium* in Rottnest Island; methane-related *Methylotenera, Methylobacter, Methylobacillus, Methylococcus* as well as biodegradation-related generalists *Geobacter, Pseudomonas, Polaromonas, Albidiferax, Burkholderia, Dechloromonas* and *Xanthomonas* in the mines cluster; thermophilic bacteria (mostly Cyanobacteria) *Roseiflexus, Synechococcus, Cyanothece, Chlorobium, Nostoc, Anabaena, Chloroflexus, Chlorobaculum* and *Synechocystis* in Little Hot Creek. LEfSe top 40 most significantly abundant functions are only found in the mines cluster, including aminoacyl-trna biosynthesis, ABC transporters, and a wide variety of amino acid metabolisms. When removing mines, most of the same functions are found in Archean Domes, except for porphyrin metabolism in Little Hot Creek and two-component regulatory system and valine, leucine and isoleucine degradation in Death Valley.

**Fig 5.**
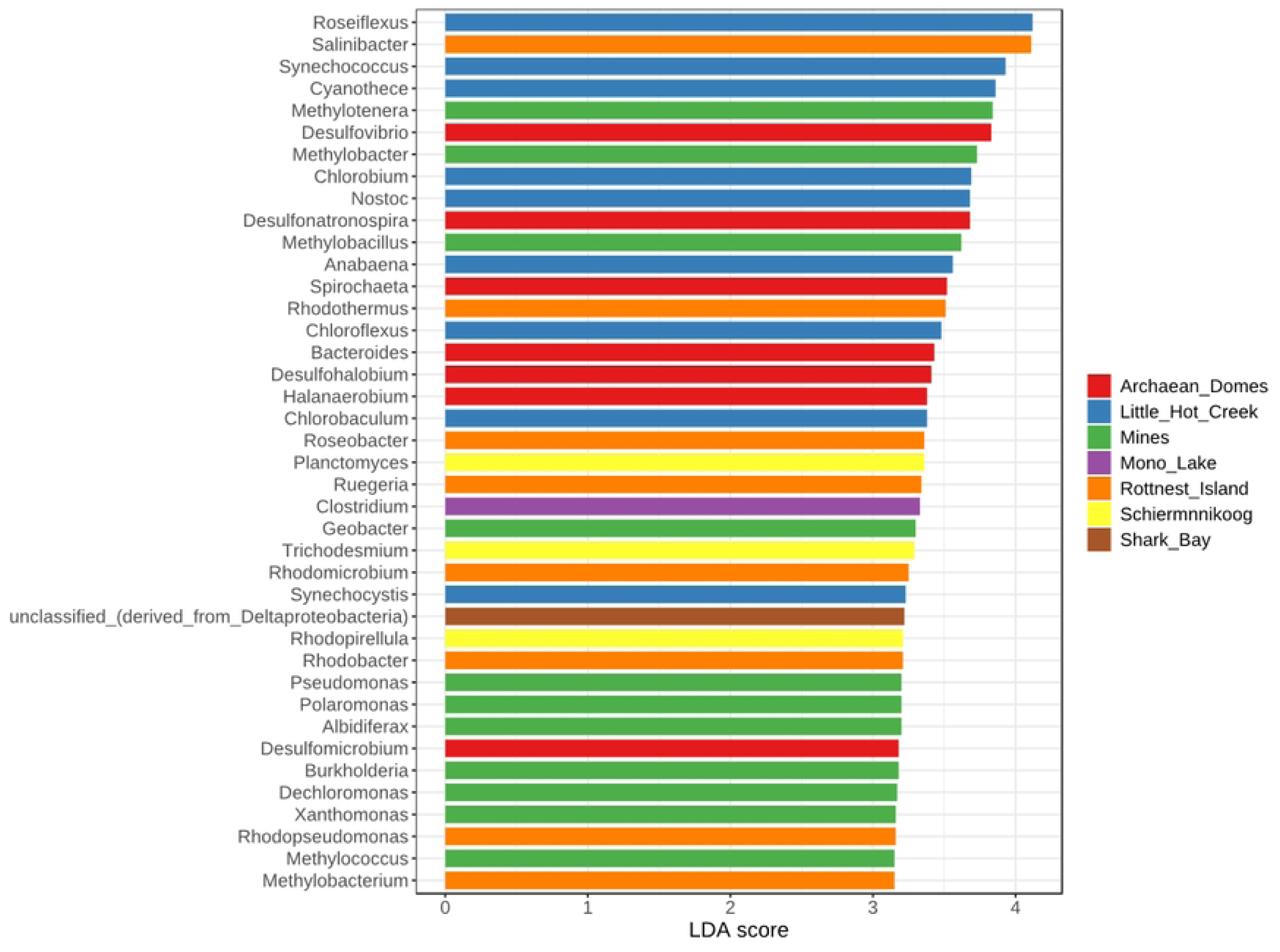
Linear discriminant analysis effect size (LEfSe) of genera. Top 40 most significant genera are depicted. The significance threshold was set to 0.01.

A total of 6 clusters of highly positively correlated top 100 most abundant bacteria in terms of relative abundances were found (Fig 6). Cluster 1 encompass mostly sulfate-reducing and green sulfur and non-sulfur bacteria, but also well-known spore-forming generalists *Bacillus* and *Clostridium*, some members of the Bacteroidaceae family (*Bacteroides, Geobacter*) and Spitochaeta. Cluster 2 includes metabolically diverse bacteria with large genomes from the Planctomycetaceae (*Planctomycetes, Pirellula, Rhodopirelilla, Blastopirellula*), Actinobacteria (*Streptomyces* and *Mycobacterium*), myxobacteria (*Anaeromyxobacter, Myxococcus, Sorangium*) and Acidobacteria (*Solibacter* and *Haliangium*) taxa. Similarly, cluster 3 includes diverse and metabolically flexible soil bacteria from the Flavobacteriaceae and Bacteroidota (mostly Cytophagales) taxa. Cluster 4 includes representatives from the purple sulfur bacteria group (*Allochromatium, Alkalilimnicola* and *Thialkalivibrio*), many metabolically diverse free-living generalists (*Shewanella, Vibrio, Pseudomonas, Burkholderia, Xanthomonas, Polaromonas, Cupravidus*) and members of Methylobacteraceae. Cluster 5 is formed exclusively by Cyanobacteria. Cluster 6 includes bacteria from 3 main groups: Rhodobacteraceae, Caulobacterales and nitrogen-fixing bacteria.

**Fig 6.**
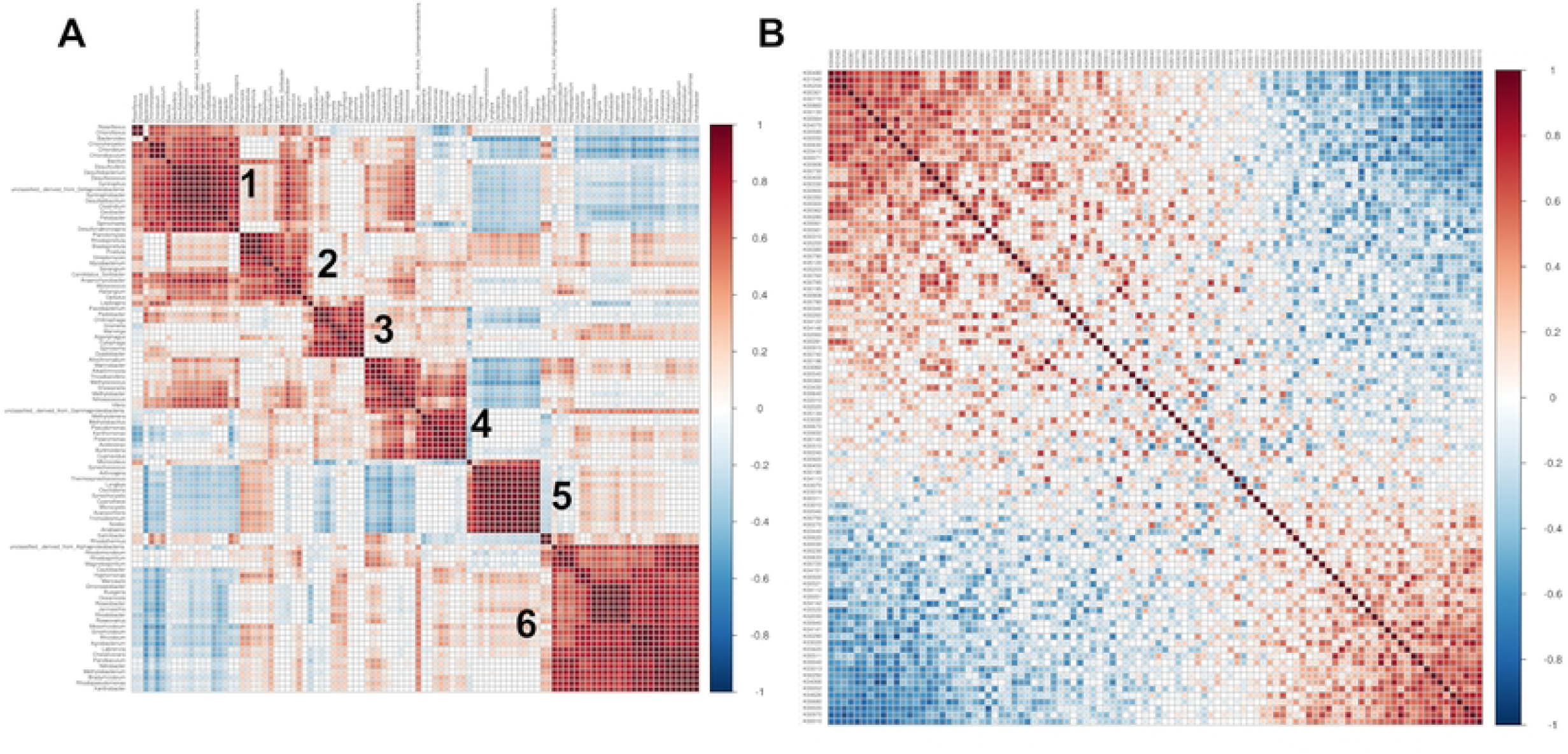
Correlation plots of relative abundances. A) Top 100 most abundant genera and B) functions are depicted. Spearman’s correlation test was used.

The correlation plot of the top 100 most abundant functions (Fig 6) shows 2 positively correlated clusters that are also negatively correlated between each other. The top left cluster contains functions related to degradation, particularly of dioxin, polycyclic aromatic hydrocarbons, naphthalene and aromatic compounds as well as microbial metabolism in diverse environments. Contrastingly, the bottom right cluster is composed of biosynthesis pathways, particularly of steroids, terpenoids and secondary metabolites. A detailed view of each function at lower hierarchical categories can be found in S3 Table.

Mantel tests of genera and functions against a compound variable of temperature-pH and a Harvesine distance matrix of geographical coordinates were performed. Of the 4 possible combinations, only the correlation between functional dissimilarity and geographic distance of microbial mats was significant (Fig 7, S4 Table).

**Fig 7.**
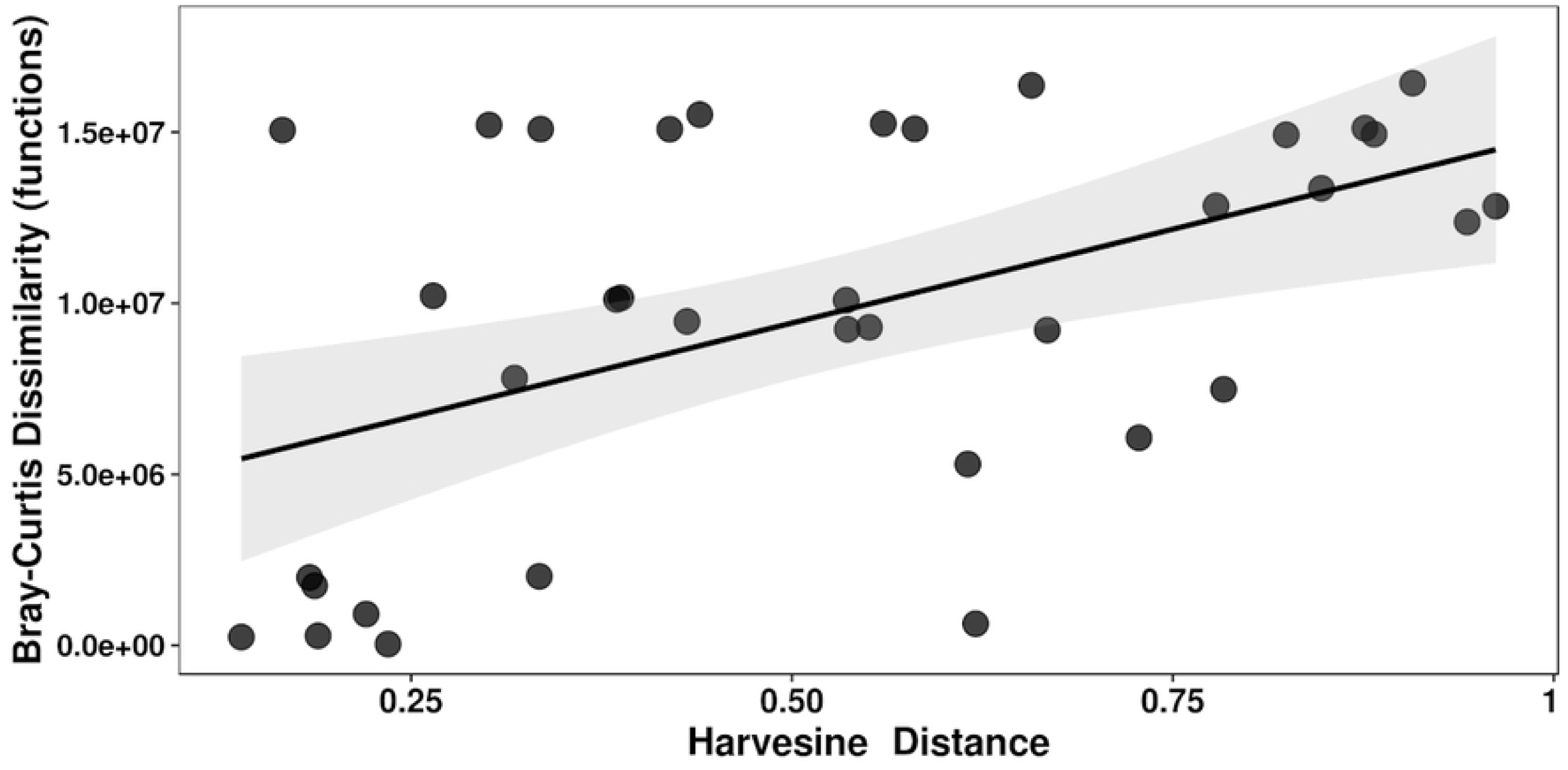
Mantel test of functions against geographic distance. A Bray Curtis dissimilarity matrix and a Harvestine distance matrix were calculated for the relative abundance of functions and the geographic distance (coordinates), respectively. The result was significant (Mantel’s r = 0.427; p = 0.0111).

## Discussion

Microbial mats are resilient and highly adaptable communities that perform mostly the same physicochemical cycles regardless of the wide range of environmental conditions they can grow at [25, 27, 28, 50]. This is explained by their functional redundancy, their remarkable nutrient scavenging and recycling abilities [32]. Thus, we expected to find similar functional and dissimilar taxonomic community compositions throughout our analysis.

We found that alpha diversity varies significantly between sites. Taxonomically, the most diverse mats were found to also be the most saline: Death Valley, Mono Lake and Shark Bay; and least diverse mat was found to be the one with higher temperature: Little Hot Creek. This aligns with previous Bolhuis et al. (2013; 2014) [51, 2] findings that salinity and fluctuating environments are some of the major drivers of microbial diversity in coastal mats. Contrastingly, Little Hot Creek being the least diverse could be explained by its high temperature, low salinity, low concentrations of sulfate and lack of seasonality which allows for fewer microclimate variations and thus, less diversity [52, 2]. We also found functional diversity to be significantly different between sites, albeit within a much narrower range than taxonomic diversity. This finding is consistent with the known high functional redundancy of microbial mats [25, 27, 28], also supporting our hypothesis; as many taxa can perform similar functions, it is expected that taxonomic diversity varies more than functional diversity across mats.

Beta diversity of both taxonomic and functional categories was also significantly different between sites. This clustering at the taxonomic but not at the functional level indicates that microbial mats show more similarities in their functional than their taxonomic composition, consistent within the functional redundancy framework. We expected to see this, as biotic interactions and metabolic codependency within each functional strata of mats would maintain similar functional composition (i.e. the Eltonian niche *sensu* Soberon, 2007 [53]) and functional redundancy would show a taxonomic structure more dependent on the general environmental conditions (i.e., the Grinellian niche niche *sensu* Soberon, 2007 [53]). Nevertheless, both alpha and beta diversity of taxonomic and functional categories still significantly differ in abundance composition within some sites. This shows not only the heterogeneity of microbial mat formations but also the challenge in spatial heterogeneity when analyzing world wide structure of microbial mats [54, 2, 51], as well as some unknown physicochemical variables and slight differences in phenotypic features within functional guilds/clusters that could drive location-dependent-differences between the analyzed clusters [55].

What defines this taxonomic and functional diversity? One explanation could be environmental filtering: structural patterns of mats being shaped by different environmental conditions. Environmental filtering refers to abiotic factors that prevent the establishment or persistence of species in a particular location [56], which captures an important process in community assembly, where species arrive at a site but fail to establish or persist due to an inability to tolerate certain abiotic conditions. And, as much as we found many expected diversity patterns that could be explained with the environments they grow in, and even though. The environmental conditions we studied do not explain our compositional differentiations, as mantel tests of temperature and pH did not seem to correlate with either taxonomic or functional composition of microbial mats (S4 Table). We consider that this observation is interesting and unexpected, given that temperature and pH are both known to influence the composition and diversity of microbial communities [57]. Miller et al (2009)[58] and Bolhuis et al (2014)[2] even refer to temperature as the environmental factor that seems to largely determine the composition of microbial mats. Still, in our study, temperature and pH do not appear to have direct influence over the compositional differences observed for world wide microbial mats. In a recent study on Archaean Domes spatial differentiation [59], the most common taxa were found to be responding to environmental variables, showing environmental filtering, while the large rare biosphere is extraordinarily diverse and does not show any sign of environmental filtering, suggesting a large “seed bank” in the deep aquifer as well as a highly dynamic composition of these mats under wet conditions. Seed banks in microbial mats could be minimizing the environmental correlation.

Having said that, considering the unavailability of more complete microenvironmental data online, the influence of environmental factors in shaping local structure of microbial mats cannot be entirely ruled out. Microbial mats are heterogeneous at the micro- and macro-scale and physicochemical gradients of light, temperature, salinity, oxygen, carbon, sulfur, and nitrogenous compounds will affect the microbial community composition at the microscale, and eventually will show large scale dissimilarities. We know from previous studies in Cuatro Cienegas that environmental conditions strongly influence the distribution of microbial mat communities by shaping metabolic niches, but the influence on taxonomic composition is rather weak, as recorded for microbiomes in marine environments [55]. Heterogeneity is typical for almost any ecosystem found to harbor microbial mat formations and is generated by alternating stable states [60], so, if not large scale abiotic dissimilarities, then local abiotic heterogeneity is where compositional correlations may lay at. This only adds difficulty in environmental data for comparison, to the already hard datasets filtering, where sufficiency of sampling efforts, same sequencing techniques for comparable studies and the ability to process data homogeneously is extremely important for eliminating as many biases as possible [61].

Nevertheless, other possible explanations could also be found to be shaping community structure besides environmental factors, such as biotic interactions. Adding to environmental filtering (*sensu* Kraft et. al., 2015 [56]), even if species are able to arrive at a site and survive in the absence of neighbors, the next ‘filtering’ logical step is to understand species abilities to persist in the presence of other interacting species already present in the community. The latter makes the founder effect crucial in understanding dissimilarities in microbial mats and microbial community composition in general [62, 63, 64, 59]. Moreover, the Mantel test (Fig 7) did show a significant moderate correlation between geographical distance and functional composition (closer mats have a more similar functional composition), pointing towards potential migration and selection of functionally compatible taxa on closer communities. Given so, and adding to them being functionally redundant systems, microbial mats would be a prime example of a taxa-function highly robust microbial community [65], however, more thorough studies aiming at this particular aspect are needed.

Similarities between microbial mats were also found in our study. At a higher hierarchy of functions (KEGG metabolism category), all microbial mats seem to behave similarly, with top 3 functional categories being nucleotide, carbohydrate and amino acid metabolisms. These results coincide with a previous metagenomic analysis of microbial mats by Santoyo (2021)[66]. However, as seen by the correlation plot of Fig 7, there are two blocks of functions at a lower hierarchy (KEGG pathways category) positively correlated within but negatively correlated between each other, one of biosynthesis and the other of degradation. Such uncorrelated contrasting blocks of functions could signal local or seasonal adaptation, or even specialization in microbial mats, particularly in the form of nutrient acquisition. This becomes more tangible when looking at the comparison between Red and Green mats from Cuatro Cienegas, which are dominated by specialist and generalist bacteria, respectively [7]. A significant differentiation in the overall metabolism of its members could lead to functional specialization of the mat like the one suggested here. In addition, the impact of generalists is of utmost importance for functional diversity and even taxonomical shaping in a given environment; microbial dispersion driven by the ability of generalist species allows for expansion across ecosystems, it means that descendants spread into new environments which makes experience distinct evolutionary pressure, and evolve into new species, whose eventually specialize to their new habitats, when a microbial guild contains high frequency of generalist species, it also tends to contain specialist species that belong to diverse environments, thus, evolutionary consequences of specialization what we are likely to see in the analyzed worldwide microbial mat [67].

Similarly to what other studies have found [26, 24], some members of the Cyanobacteria group were found to be highly prevalent in microbial mats across the globe. For hypersaline mats, Cyanobacteria has been found to be the dominant primary producers and are responsible for the production of a thick exopolysaccharide matrix that serves among others as protection against desiccation [2]. In our study, we found *Cyanothece* to be particularly important, followed by *Microcoleus. Cyanothece* was previously found to be an abundant member of hypersaline mats from Chile [68] and thermopfilic mats from East Asia [69], while *Microcoleus* has been deemed a mat-building Cyanobacteria by Moezelaar et al (1996)[70] and also found as a dominant species in hypersaline mats from Guerrero Negro [71], eutrophic marine mats [72, 73], alkaliphilic and halophilic mats [74]. Our results further support the role of these taxa as keystone species in microbial mats irrespective of the environmental conditions. Additionally, we found 2 Planctomycetes genera, *Planctomyces* and *Rhodopirellula*, also to be highly prevalent. Fernandez et al., (2016)[75] found Planctomycetes in the bottom layer of hypersaline mats from Chile, and suggested they play a key structural role. Also, Rozanov et al (2017)[76] found Planctomycetes to be highly abundant in the middle layers of thermophilic mats. Santoyo (2021)[66] found Planctomycetes in microbial mats from contrasting environments as well. Furthermore, *Planctomyces* has been found to be a dominant species in mats from hydrothermal vents [77] and *Rhodopirellula* in algal mats [78]. However, this is the first record of the Planctomycetes group as a potential keystone taxon for microbial mats.

Clusters from the correlation plot (Fig 7) show groups of bacteria that are often found together across microbial mats. Cluster 1 mostly includes sulfate-reducing, green sulfur and non-sulfur bacteria and spore-forming generalists. Sulfur-related bacteria are well-known members of microbial mats, perpetuating necessary environmental gradients to keep the mat functioning [2]. However, there is no previous evidence of interaction or common environmental preferences between them and *Bacillus* or *Clostridium* other than high temperature tolerance [79,80]. Clusters 2, 3 and 4 include metabolically flexible, generalist bacteria with large genomes. Generalist bacteria have been found to increase resilience of bacterial communities [81] and, although often overlooked in microbial mats, they may play an important role by providing genetic redundancy and alternative metabolic pathways [82]. Noteworthy, *Planctomyces* and *Rhodopirellula* appear in cluster 2, further supporting their potential role as keystone species. Cluster 5 is entirely composed of Cyanobacteria. Due to the principle of competitive exclusion [83], one would not expect a strong positive correlation between ecologically equivalent keystone members of a community. Previous studies have shown competitive exclusion between Cyanobacteria to be common in aquatic systems [84, 85, 86, 87]. Nonetheless, Litchman et al. (2003)[88] found that fluctuations in light supply, including daily light-dark cycles, may lead to a long-term persistence of more than 1 species of cyanobacteria. This could be explained by Cyanobacteria’s wide vertical distribution, inherent to their ability to migrate along light and chemical gradients on the different mat layers [54, 89, 90, 91, 92]. In general, it is known that spatial heterogeneity and temporal variation of the environment help to prevent competitive exclusion [93], both of which are intrinsic characteristics of microbial mats and their environment. Moreover, niche partitioning, which is likely to play a role in allowing ecologically similar members of microbial mats to coexist in general [94, 95], has been observed at the phototrophic fraction of microbial mats [96]. Although Cyanobacteria may be ecologically highly similar, fluctuating environmental conditions coupled with niche partitioning may allow them to coexist in the upper layers of microbial mats. Cluster 6 encompasses mostly nitrogen-fixing and halophilic bacteria. It is known that nitrification and denitrification are likely general phenomena for hypersaline mats due to the favorable conditions therein [97] and that microorganisms in hypersaline environments contribute to the biogeochemical cycles involving carbon and nitrogen as sources of energy [98]. Thus, our results add to previous findings of an increased importance of nitrogen metabolism in hypersaline mats [66].

Although correlations between bacterial abundance were found across mats, some bacteria tend to be overly abundant in certain sites. For instance, Cluster 5 (Cyanobacteria) is significantly more abundant in Little Hot Creek and Cluster 6 (halophiles and nitrogen bacteria) in Rottnest Island. Moreover, specific groups of bacteria are found as significantly more abundant in specific sites, coherent with their ecological roles and the environmental conditions of each site. Physicochemical conditions may serve as an initial environmental filter [99] from which mats establish and certain taxa draw competitive advantages. Even though we did not find a significant correlation between temperature and pH nor geographical distance and beta diversity, this result by itself may suggest a predominance of Grinellian niche dynamics are more important for the most abundant taxa in microbial mat community assembly [53]. Even though, the analysis of rare taxa as previously done in Archaean Domes may present a more Eltonian niche dynamics, where through a phylogenetic diversity and dispersion analyses, it showed that the rare biosphere is subject to dispersal and drift more than abiotic filtering or competitive exclusion [59].

## Acknowledgments

Mariette Viladomat Jasso is a doctoral student from Programa de Doctorado en Ciencias Biomédicas, Universidad Nacional Autónoma de México (UNAM), and has received CONACyT fellowship 736510. We want to thank Laura Espinosa-Asuar and Erika Aguirre-Planter for technical assistance, as well as Rodrigo Zorrilla for his help during fieldwork. The authors declared no potential conflicts of interest and have approved the final version of the manuscript.

## Supplementary information

**S1 Fig. Rarefaction curves of metagenomes**.

**S2 Fig. Core analysis of genera and functions**. Minimum prevalence was set to 20% for genera (A) and 99% for functions (B) for easier visualization.

**S1 Table. Metagenomic sampling and corresponding publications**.

**S2 Table. Environmental measurements of metagenomes**.

**S3 Table. KEGG accession numbers of functional categories**.

**S4 Table. Results of Mantel tests**.

